# Tree size drives diversity and community structure of microbial communities on the bark of beech (Fagus sylvatica)

**DOI:** 10.1101/2022.01.19.476951

**Authors:** Lukas Dreyling, Imke Schmitt, Francesco Dal Grande

## Abstract

Tree bark constitutes ideal habitat for microbial communities, because it is a stable substrate, rich in micro-niches. Bacteria, fungi, and terrestrial microalgae together form microbial communities, which in turn support more bark-associated organisms, such as mosses, lichens, and invertebrates, thus contributing to forest biodiversity. We have a limited understanding of the diversity and biotic interactions of the bark-associated microbiome, as investigations have mainly focussed on agriculturally relevant systems and on single taxonomic groups. Here we implemented a multi-kingdom metabarcoding approach to analyse diversity and community structure of the green algal, bacterial, and fungal components of the bark-associated microbial communities of beech, the most common broadleaved tree of Central European forests. We identified the most abundant taxa, hub taxa, and co-occurring taxa. We found that tree size (as a proxy for age) is an important driver of community assembly, suggesting that environmental filtering leads to less diverse fungal and algal communities over time. Conversely, forest management intensity had negligible effects on microbial communities on bark. Our study suggests the presence of undescribed, yet ecologically meaningful taxa, especially in the fungi, and highlights the importance of bark surfaces as a reservoir of microbial diversity. Our results constitute a first, essential step towards an integrated framework for understanding microbial community assembly processes on bark surfaces, an understudied habitat and neglected component of terrestrial biodiversity. Finally, we propose a cost-effective sampling strategy to study bark-associated microbial communities across large spatial or environmental scales.

## Introduction

The aboveground surfaces of plants are ideal substrates for microbial colonization. The bark surface (or dermosphere; Lambais et al., (2014)), in particular, is one of such important aboveground substrates in forests. The bark provides a range of microhabitats that promote colonization of microbial communities with varied ecologies (Whitmore, 1963). On the one hand, microsites such as holes, cracks, and lenticels retain humidity and nutrients, thus constituting stable microhabitats suitable for slow-growing, stress-sensitive microbes. On the other hand, the exposed surfaces of the bark may harbour more stress-resistant microbial communities that can cope with environmental challenges (Vorholt, 2012; Aguirre-von-Wobeser et al., 2021), such as low nutrient availability, increased exposure to light, fluctuating moisture conditions and desiccation (Lindow and Brandl, 2003; Vorholt, 2012; Leff et al., 2015), and presence of compounds that are resistant to microbial degradation (e.g., suberin), or that directly inhibit microbial growth (Baldrian, 2017).

Compared to other aboveground components, such as leafs, branches or fruits, that undergo seasonal and diurnal changes (Vitulo et al., 2019), bark represents a stable, long-lived substrate that supports microbial colonization (Leff et al., 2015). Further, the bark surface is often screened from excessive precipitation- and/or UV radiation by the tree canopy and changes slowly during development over several years (Whitmore, 1963). A number of studies have investigated the bark-associated microbial diversity, especially for fungi and bacteria, in various systems, e.g., grapevine plants (Martins et al., 2013; Arrigoni et al., 2018), bark beetle-infested spruce (Strid et al., 2014), *Ginkgo* (Leff et al., 2015) and avocado trees (Aguirre-von-Wobeser et al., 2021). These studies report that the tree bark supports microbial communities that are often distinct from spatially-close substrates like leaves and roots (Martins et al., 2013; Leff et al., 2015; Arrigoni et al., 2018), indicating niche differentiation and a clearly structured habitat (Aguirre-von-Wobeser et al., 2021). Furthermore, the dermosphere constitutes a reservoir for microbial diversity (Arrigoni et al., 2018; Hagge et al., 2019; Kobayashi and Aoyagi, 2019), potentially harbouring undiscovered specialist taxa (Aschenbrenner et al., 2017), and taxa that facilitate the colonization of other epiphytes, including lichens (Aschenbrenner et al., 2017). The microbial communities on tree bark, and the biofilm which they form, can indeed be considered the basis of a food web that supports photosynthetic epiphytes (e.g., mosses and lichens), as well as a diverse microfauna (Andre, 1985). With an estimated more than 3 trillion trees in the world (Crowther et al., 2015), bark communities could thus be particularly important reservoirs of biological diversity. However, bark is a poorly explored habitat with respect to microbial diversity and community structure, compared to other substrates such as the phyllosphere and rhizosphere.

Biological and environmental factors driving diversity and community assembly in bark-associated epiphytes have been linked to forestry management, e.g., management intensity (Boch et al., 2021), forest homogeneity (Lamit et al., 2015), deadwood abundance (Boch et al., 2021), and tree age. For the latter, higher epiphyte diversities have been linked to the availability of large, old-growth trees (Aude and Poulsen, 2000; Nascimbene et al., 2013; Boch et al., 2021), probably because of higher niche partitioning in older trees (Łubek et al., 2020). At smaller spatial scales, abiotic drivers of bark-associated diversity and community structure include ultraviolet radiation, water shortages and correlated desiccation, poor nutrient availability (Lindow and Brandl, 2003; Vorholt, 2012; Leff et al., 2015), while biotic drivers include host traits, such as maturity of the substrate and host genotype (Arrigoni et al., 2018, 2020). Community composition is therefore, to some degree, host specific. A few studies showed that the trends observed for macroepiphytes (e.g., bryophytes, lichens) or components of the phyllosphere also appear in bark-associated microbes (e.g., Vorholt, 2012; Arrigoni et al., 2020). However, our understanding of the factors shaping the different components of the highly diverse bark-associated microbial communities is still limited.

Most of the studies focus on non-natural, commercially driven ecosystems like orchards or vineyards (Martins et al., 2013; Arrigoni et al., 2018). Studies are often conducted over small spatial scales with small sample sizes (e.g., Leff et al., 2015). Lastly, the focus often lies on only a single group of microorganisms, with bacteria and fungi far outweighing terrestrial algae (Aschenbrenner et al., 2017; Petrolli et al., 2021). Integrative sampling of major microbial contributors over regional or potentially even global scales can help identifying not only the diversity of microorganisms but also potential cooperative and competitive interactions among them. Revealing the diversity and structure of these rather unique microbial communities is essential to predict their responses in a changing environment. Furthermore, considering the importance of fungi, bacteria and algae to ecosystem nutrient and energy budgets in terrestrial habitats, gaining information on the bark-associated microbial communities and their dynamics is essential and directly relevant for ecosystem service assessment.

In this study we present one of the first integrated investigations of the bark microbiome in temperate forests. Here we use the term microbiome following the definition by Berg et al., (2020). Specifically, we study the three main components of the bark microbiome, i.e. green algae, bacteria and fungi. We sampled bark surfaces in forests under different management regimes, ranging from highly-managed stands to relatively undisturbed sites in the core zone of a national park. We used metabarcoding to analyse microbial diversity, community structure and species interactions from the tree to the landscape level. Specifically, we asked the following questions: i) What is the microbial diversity found on the bark of the most common broadleaved tree in central Europe (*Fagus sylvatica*)?, ii) Which species co-occur and who are the main players in the identified ecological modules?, iii) Which factors, i.e. management intensity and tree-size classes (as a proxy for tree age), affect the bark-associated microbiome, both at tree and landscape level?

The comparison of diversities among trees of different size classes within a spatially-explicit framework allowed us to test for the effects of sampling design on the estimation of microbial diversity. This information is essential for further, larger scale sampling campaigns.

## Material & Methods

### Study site and sampling

Sampling sites are situated within the central European region of Hainich-Dün (Thuringia, Germany), that is one of the three regions of the Biodiversity Exploratories project (Fischer et al., 2010). The Hainich-Dün region is characterized by soils stemming from calcareous bedrock, an annual rainfall between 500-800 mm, a mean temperature of 6.5-8 °C at an elevation of 285-550 m above sea level (Fischer et al., 2010).

Sample collection took place in autumn between the 13^th^ and 15^th^ of October 2020. We chose a subset of 16 out of the established 50 experimental plots (Fischer et al., 2010), sampling a subplot of 20×20 m within the 100×100 m experimental plots. These plots were chosen to represent two regimes of land-use intensity, namely a high and a low intensity forest management (eight plots each), according to the Forest Management Index (ForMI, high > 1, low < 1). This is an index combining measures of harvested stem volume, occurrence of non-natural species and deadwood stemming from harvest (Kahl and Bauhus, 2014). Specifically, we defined three size classes: large (i.e. >30 cm diameter at breast height (DBH)), medium (15 – 30cm DBH) and small (5 – 15cm DBH). Two trees per size class were sampled. When one tree-size class was not available (three plots), we sampled more trees of the other size classes depending on which was highly abundant in the direct vicinity as judged in the field. Within each plot we recorded the spatial position of the trees relative to each other by measuring distance (m) and azimuth (degrees) from the nearest sampled tree.

We collected microbial bark surface communities using individually wrapped sterile nylon-flocked medical swabs with a 30 mm breakpoint, typically used for medical specimen collection (FLOQSwabs™, Copan, Brescia, Italy). The breakpoint mechanism minimises the possibility of contamination when transferring the swab into the Eppendorf tube. Prior to collection the bark was moisturised with deionized water to mobilize the surface biofilms. Then the tree was swabbed at breast height in a 3 cm-wide band around the trunk, rolling the swab and moving it up and down while applying gentle pressure. While swabbing, we took care to include smooth surfaces as well as cracks and crevices in the bark, to ensure a good representation of micro-habitats. The swab head was broken off into Eppendorf tubes pre-filled with 750 μl Nucleic Acid Preservation (NAP) buffer (Camacho-Sanchez et al., 2013). Tubes were immediately placed in styrofoam boxes with ice and the samples were subsequently stored at 4 °C until DNA extraction.

### DNA extraction

Prior to DNA extraction we added 750 μl ice-cold phosphate-buffered saline (PBS) into the Eppendorf tube and centrifuged the sample for 15 min at 6000×g as recommended by Menke et al., (2017). The supernatant was then discarded without disturbing the swab head or pellet. DNA was extracted using the Quick-DNA Fecal/Soil Microbe Microprep kit (Zymo Research Europe GmbH, Freiburg). Initial tissue lysis was achieved through mechanical disruption by bead beating, using the beads included in the extraction kit. We modified the kit protocol by directly adding the beads and bead-beating buffer into the tube containing the pellet and swab and shaking for a total of 6 min (SpeedMill PLUS, Analytik Jena, Jena, Germany). In the later steps we followed the manufacturer’s protocol, using DNAse-free water as elution buffer. We included six contamination controls, that were sequenced as well. DNA extracts were frozen at −20 °C until PCR.

### PCR amplification and high-throughput sequencing

Algal, bacterial and fungal fractions of the extracted microbial DNA were amplified, using primers for the ITS2 region for fungi and algae, and the 16S hyper-variable region V3 – V4 for bacteria (Table 1).

**Table 1:**
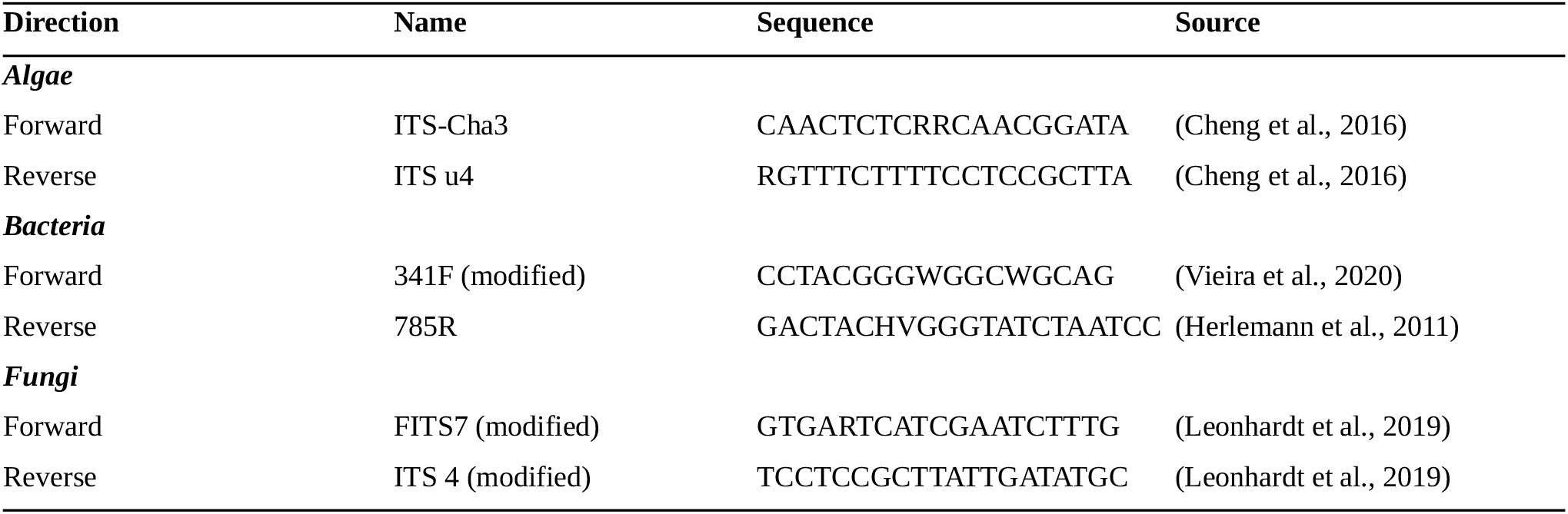
Primer names and sequences used in this study.

All samples were amplified in duplicate with forward and reverse primers individually tagged with octamers allowing for a double index multiplexing approach. Every replicate contained eight negative controls (i.e., master mix without sample), as well as 16 “Multiplex Controls” (i.e., empty wells) to allow detection of potential primer jump during sequencing (Schnell et al., 2015). We set up 15 μl PCR reactions containing 5 ng of DNA, 7.5 μl of MyTaq™ HS Mix, 2x (Bioline GmbH, Luckenwalde, Germany), 0.6 μl 10 μM of each primer, and 4.3 μl DNAse free water.

Cycling conditions differed in cycle number and annealing temperature among organismal groups. Conditions were as follows: an initial denaturation at 95 °C for one minute, followed by 30 (algae, bacteria) or 35 (fungi) cycles of denaturation at 95 °C for 15 s, annealing at 54 °C (algae), 59 °C (bacteria) or 56 °C (fungi) for 15 s and elongation at 72 °C for 10 s, with a final extension at 72 °C for 1 min. Samples were randomly distributed over two 96-well plates, with both replicates following the same placement scheme.

The amplicons were individually cleaned using magnetic beads (MagSI-NGS^Prep^ Plus, magtivio B.V., Geelen, Netherlands) and DNA concentration was quantified with fluorescence measurement using the Qubit dsDNA HS assay (Thermo Fisher Scientific, MA, USA) as specified by the manufacturer. The replicates were equimolarly pooled within the respective organismal groups, creating a total of three pools for sequencing. The pooled amplicons were send for library preparation and sequencing to Fasteris SA (Plan-les-Ouates, Switzerland). Libraries were prepared for each pool according to the Fasteris MetaFast protocol (https://www.fasteris.com/), in order to avoid PCR for library preparation and thus minimizing additional PCR bias and chimera creation. The samples were sequenced on an Illumina MiSeq (Illumina Inc., San Diego, CA, USA) with 2×300 bp paired-end reads.

### Bioinformatics

Adapter-trimmed reads trimmed with Trimmomatic (Bolger et al., 2014) were supplied by the sequencing provider. We demultiplexed the reads using Cutadapt v3.3 (Martin, 2011) following the demultiplexing combinatorial dual-indexes section of the manual. The error rate was set to 0.15, allowing no insertions or deletions, and discarding reads shorter than 50 bp. Commands were run a second time with the octamer tags in the reverse order to account for amplicons in mixed orientation resulting from PCR-free library preparation. The resulting files were merged to obtain one R1 and one R2 file per replicate. Reads were checked for remaining primer sequences, which were removed using Cutadapt, if present.

The demultiplexed reads were further processed with the DADA2 pipeline (Callahan et al., 2016). Filtering and trimming operations used DADA2 default parameters, except for setting a truncation length (truncLen = c(250,260)) for bacteria, but not for algae and fungi since the length of the ITS2 region can vary between taxa (Schoch et al., 2014). Furthermore, the maximum error rates were relaxed to maxEE = c(5,5) for bacteria and maxEE = c(6,6) for algae and fungi. After de-noising and sample inference, pairs were merged within each replicate, chimeras were removed and one amplicon sequence variant (ASV) table was constructed per replicate. To account for the mixed orientation of the libraries we checked the tables for reverse complement sequences, reversed them and added their counts to the respective complement sequence using DADA2s *rc()* function.

Finally, the replicates were merged by summing up the read counts.

For taxonomic assignment the sequences were matched against publicly available databases, namely UNITE general release 8.2 (Abarenkov et al., 2020) for fungal reads and SILVA 138.1 SSU Ref NR 99 (Quast et al., 2012) for bacteria. Since no similar database is currently available for green algae we used the program SEED2 v2.1.2 (Větrovský et al., 2018) to conduct a BLASTn search against GenBank (Clark et al., 2016).

We then checked the reads for potential contamination with the *decontam* package (Davis et al., 2018), using the combined prevalence and frequency approach. For all organismal groups d*econtam* only showed low numbers (algae = 0, bacteria = 8, fungi = 4) of potential contaminant ASVs, which were discarded. The *decontam-*filtered ASV tables were curated using the LULU algorithm (Frøslev et al., 2017) to merge highly similar ASVs and obtain more reliable diversity metrics. Taxonomic information for all ASVs can be found in Supplementary Table 1.

### Diversity and community structure analyses

All analyses were conducted in R (R Core Team, 2021, version 4.0.4) through RStudio (RStudio Team, 2021). ASV tables, taxonomic information and accompanying metadata were combined using the *phyloseq* R package (McMurdie and Holmes, 2013) to ease analyses. Figures were created with *ggplot2* (Wickham, 2016) and *gridExtra* (Auguie, 2017). Samples were not rarefied as recommended by McMurdie and Holmes (2014) and instead treated as compositional count data (Gloor et al., 2017). Script of all analyses are available on GitHub at https://github.com/LukDrey/beech_micro_communities. [link active upon publication; the code is made available to Reviewers as supplementary file].

### Intra-group diversities

We calculated the Shannon Index (Shannon, 1948) as a measure of alpha diversity, using the function *estimate_richness()* on the full untransformed ASV table as obtained from DADA2 and LULU. Differences in Shannon diversity between tree sizes and management category were tested with an Analysis of Variance (ANOVA) with the function *aov()* and verified via a Tukey Honest Significant Differences test (Tukey HSD). Furthermore, we tested whether the Shannon diversity for trees within a plot was spatially autocorrelated. For this purpose, we computed Moran’s I (method from Gittleman and Kot, 1990) as a measure of spatial autocorrelation with the function *Moran*.*I* from the ape R package (Paradis and Schliep, 2019).

To create the community barplots we aggregated taxa at the order rank and subset the datasets to the 25 relatively most abundant taxa with *get_top_taxa()* (Teunisse, 2017). The resulting subsets were transformed to reflect their compositional nature by *transform()* and plotted using *plot_composition()*, both from the *microbiome* R package (Lahti and Shetty, 2017).

### Inter-group differences

Before comparing differences in community composition of differently sized trees and management regimes, each full dataset was transformed based on centred log-ratios (CLR) with *transform()*. After the transformation we conducted a principal component analysis (PCA) on the clr-transformed datasets using the *phyloseq* function *ordinate()* (“RDA” method) which for clr-transformed data is the same as PCA. In the ordination plots we show the first two Principal Components (PC), with the axes scaled to the proportion of variance the PC explains, as recommended by Nguyen and Holmes (2019).

To test if groups showed similar within-group variance, we used the *betadisper()* function and verified the results with the accompanying permutation test *permutest()* from the *vegan* package (Oksanen et al., 2020). To test for differences in community composition between tree sizes and management intensity we performed Permutational Analysis of Variance (PERMANOVA) with a distance matrix based on Aitchison’s distance (method = “euclidean” with the *phyloseq* function *distance()* for clr-transformed data). The PERMANOVA was computed using the *vegan* function *adonis2()* examining marginal effects of tree size and management intensity together.

### Species interactions

The interaction networks were generated with the *SPIEC-EASI* method (Kurtz et al., 2015), a robust method for the sparse and compositional nature of microbiome datasets implemented in the R package *SpiecEasi*. Before network inference the ASV tables were subset to contain only ASVs contributing at least one percent of the total reads to ease both visualization and computational load. The main SPIEC-EASI algorithm was set to use the meinshausen-bühlmann’s neighborhood selection (Meinshausen and Bühlmann, 2006) and Bounded StARS model selection (Müller et al., 2016) on 50 subsamples (rep.num = 50), with lambda.min.ratio = 0.1, nlambda = 100, pulsar.select = TRUE and seed =10010. We calculated one network per organismal group, as well as one containing all three groups together.

The obtained models were refit, turned into *igraph* (Csardi and Nepusz, 2006) objects and loaded in *Gephi* v0.9.2 (Bastian et al., 2009) for visualisation. Modularity and betweenness centrality (for visualisation purposes) were computed with *Gephi*s internal algorithms (Brandes, 2001; Blondel et al., 2008). The graph layouts were constructed using the Fruchterman-Reingold algorithm (Fruchterman and Reingold, 1991). For each network, hub taxa were calculated based on vertex betweenness centrality using the *igraph* function *betweeness()* with default parameters, except setting directed = FALSE. The top five hub taxa, based on vertex betweenness centrality, were extracted.

### Differential abundance analysis

Differential abundance analysis was conducted using *ALDEx2* (Fernandes et al., 2013, 2014; Gloor et al., 2016). We compared abundances of two groups, i.e. high/low management intensity, large/small, large/medium and medium/small trees, for each organismal group. *ALDEx2* generates Monte Carlo samples (N = 128), drawn from the Dirichlet distribution for each individual sample, and tests differences between specified groups through Wilcoxon rank-sum tests. *ALDEx2* is a robust choice for compositional datasets because the data is clr-transformed internally. Taxa were declared differentially abundant if they showed a Benjamini-Hochberg corrected p-value < 0.05.

## Results

### Intra-group diversities

In total we obtained on average 38,223 reads per sample for algae (min = 12,913, max = 87,526), 58,670 reads for bacteria (min = 223, max = 181,567) and 44,239 reads for fungi (min = 9,803, max = 117,957), with minima of 93, 103 and 92 reads and maxima of 16,436, 19,798 and 76,422 reads for the sequencing controls. From these reads, we retrieved 216 algal, 1,723 bacterial and 992 fungal ASVs. Overall algae and fungi displayed similar Shannon alpha diversity, while bacteria showed a slightly higher diversity (Figure 1). Neither algae, fungi nor bacteria exhibited statistically significant differences in alpha diversity when comparing low and high management intensity plots (Figure 1 (A), (B) and (C)). Considering differences between tree sizes, smaller trees displayed higher alpha diversity for algae and fungi (Figure 1 (D) and (F)). Overall, bacterial diversities were more uniform, but displayed higher median Shannon diversity values for larger trees (Figure 1 (E)). This trend was corroborated by the results of an ANOVA comparing the three tree-size classes. For algae, we found a significant overall effect of tree size (F = 4.163, p < 0.05), that was driven by a significant difference between large and small trees (post-hoc test p-value < 0.05). No significant differences were found between large/medium and medium/small trees. Tree size had a significant effect overall (F = 16.499, p < 0.001) in fungi, with significant differences between large and medium (p < 0.01), as well as large and small trees (p < 0.001). For bacteria we found no significant overall effect.

**Figure 1.**
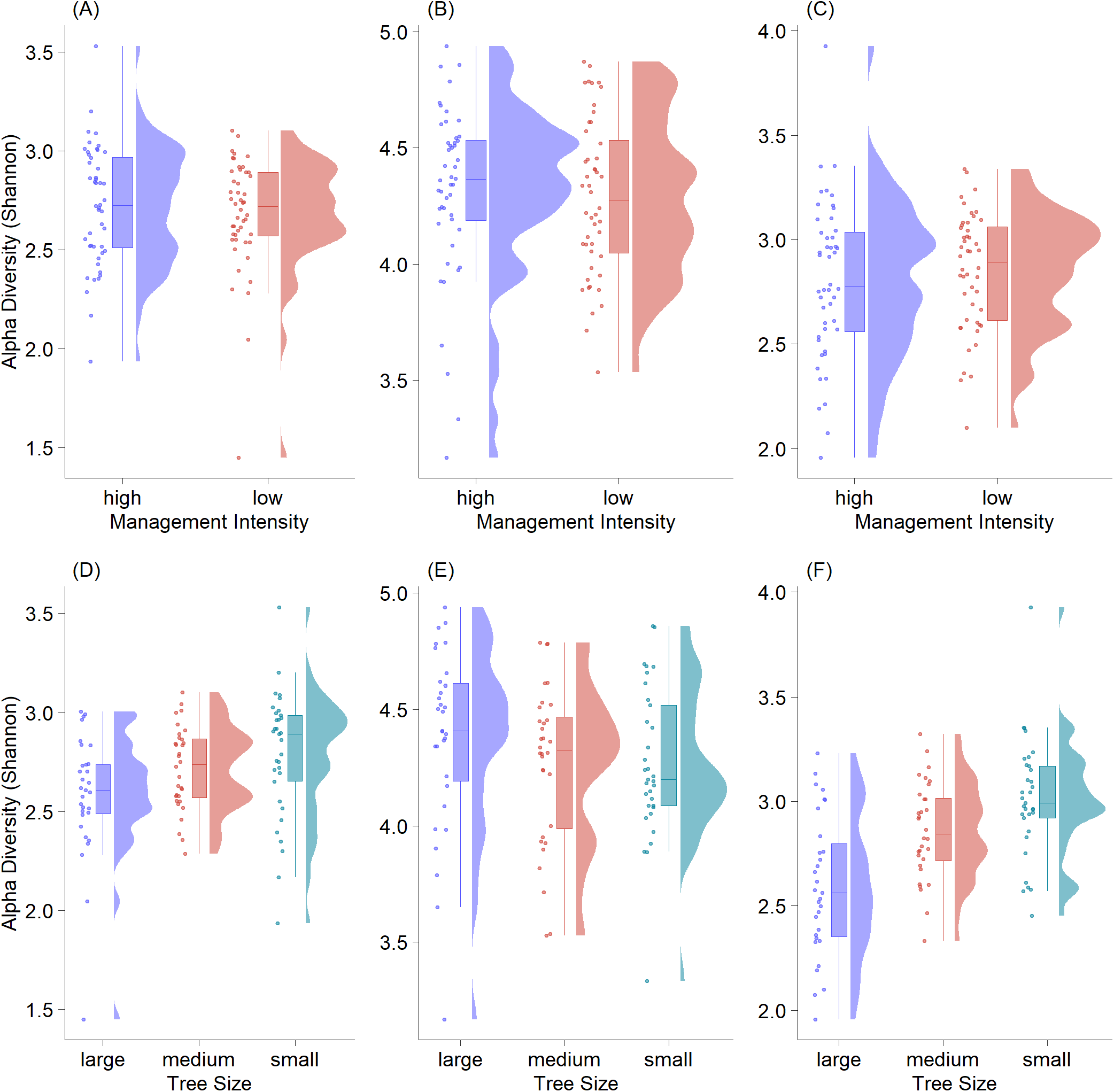
Rain-Cloud plots of alpha diversity (Shannon) against management regime and tree size for algae (A and D), bacteria (B and E) and fungi (C and F). Differences are visible from the boxplots, while the original data structure is visible from raw data scatters (randomly jittered) and raw data distribution.

Spatial autocorrelation tests showed that only in four of 48 cases the null-hypothesis of no spatial correlation could be rejected (Table 2). This indicates that the effect of spatial autocorrelation within plots is negligible. Trees belonging to the same plot showed very similar Shannon alpha diversity values. One exception is plot HEW8 where alpha diversity was significantly spatially autocorrelated for both algae and bacteria.

**Table 2.** Overview of Moran’s I (MI) values for all three organismal groups in the studied plots. An observed higher significant p-value (< 0.05) vs. expected indicates positive autocorrelation, whereas lower MI values indicate negative autocorrelation.

| Algae |  |  |  | Bacteria |  |  |  | Fungi |  |  |  |  |
| --- | --- | --- | --- | --- | --- | --- | --- | --- | --- | --- | --- | --- |
| Plot ID | MI | MI | sd | P value | MI | MI | sd | P value | MI | MI | sd | P value |
|  | (observed) | (expected) |  |  | (observed) | (expected) |  |  | (observed) | (expected) |  |  |
| HEW11 | -0.19 | -0.2 | 0.17 | 0.95 (n.s.) | -0.09 | -0.2 | 0.12 | 0.32 (n.s.) | 0 | -0.2 | 0.24 | 0.41 (n.s.) |
| HEW20 | -0.41 | -0.2 | 0.16 | 0.17 (n.s.) | -0.15 | -0.2 | 0.14 | 0.72 (n.s.) | -0.43 | -0.2 | 0.16 | 0.13 (n.s.) |
| HEW26 | -0.26 | -0.2 | 0.24 | 0.8 (n.s.) | -0.55 | -0.2 | 0.21 | 0.09 (n.s.) | -0.24 | -0.2 | 0.23 | 0.86 (n.s.) |
| HEW27 | -0.46 | -0.2 | 0.2 | 0.19 (n.s.) | -0.09 | -0.2 | 0.19 | 0.54 (n.s.) | -0.27 | -0.2 | 0.24 | 0.77 (n.s.) |
| HEW28 | 0.32 | -0.2 | 0.3 | 0.09 (n.s.) | -0.33 | -0.2 | 0.22 | 0.55 (n.s.) | -0.39 | -0.2 | 0.31 | 0.53 (n.s.) |
| HEW31 | -0.09 | -0.2 | 0.11 | 0.31 (n.s.) | -0.36 | -0.2 | 0.16 | 0.31 (n.s.) | 0.02 | -0.2 | 0.14 | 0.12 (n.s.) |
| HEW32 | 0.04 | -0.2 | 0.13 | 0.06 (n.s.) | -0.18 | -0.2 | 0.09 | 0.84 (n.s.) | 0.05 | -0.2 | 0.15 | 0.09 (n.s.) |
| HEW33 | -0.22 | -0.2 | 0.25 | 0.93 (n.s.) | -0.21 | -0.2 | 0.08 | 0.9 (n.s.) | 0.04 | -0.2 | 0.29 | 0.41 (n.s.) |
| HEW34 | -0.3 | -0.2 | 0.16 | 0.52 (n.s.) | -0.3 | -0.2 | 0.19 | 0.62 (n.s.) | -0.35 | -0.2 | 0.2 | 0.47 (n.s.) |
| HEW35 | -0.35 | -0.2 | 0.14 | 0.29 (n.s.) | -0.14 | -0.2 | 0.15 | 0.71 (n.s.) | 0.09 | -0.2 | 0.13 | 0.03 (*) |
| HEW36 | -0.31 | -0.2 | 0.12 | 0.39 (n.s.) | -0.2 | -0.2 | 0.11 | 0.99 (n.s.) | -0.12 | -0.2 | 0.09 | 0.37 (n.s.) |
| HEW37 | 0.01 | -0.2 | 0.11 | 0.07 (n.s.) | -0.2 | -0.2 | 0.09 | 0.98 (n.s.) | -0.27 | -0.2 | 0.11 | 0.53 (n.s.) |
| HEW43 | -0.31 | -0.2 | 0.15 | 0.48 (n.s.) | -0.34 | -0.2 | 0.09 | 0.14 (n.s.) | -0.34 | -0.2 | 0.08 | 0.08 (n.s.) |
| HEW49 | -0.15 | -0.2 | 0.1 | 0.58 (n.s.) | -0.38 | -0.2 | 0.11 | 0.1 (n.s.) | -0.13 | -0.2 | 0.08 | 0.44 (n.s.) |
| HEW5 | -0.06 | -0.2 | 0.09 | 0.09 (n.s.) | -0.29 | -0.2 | 0.1 | 0.38 (n.s.) | -0.01 | -0.2 | 0.09 | 0.04 (*) |
| HEW8 | 0.31 | -0.2 | 0.22 | 0.02 (*) | 0.38 | -0.2 | 0.26 | 0.02 (*) | 0.25 | -0.2 | 0.28 | 0.11 (n.s.) |

Bark microbial communities were similar among trees, with only minor differences in rare orders for all three organismal groups. For algae, the predominant orders were Trebouxiales and Chlorellales, with Trebouxiales contributing more than 50% of the reads in many plots (Figure 2 (A)). Rare orders displayed a relatively high inter-plot variability, with Prasiolales being almost absent for the plot HEW5. Compared to algae, Bacteria were more homogeneous, with the same four orders - Rhizobiales, Sphingomonadales, Acetobacterales and Cytophagales - displaying comparably high relative abundances in all plots (Figure 2 (B)). We found a higher rare-order diversity in bacteria and fungi compared to algae. Capnodiales were by far the most abundant fungal order, and dominant in all plots (Figure 2 (C)). Compared to bacteria and algae, more fungal reads could not be assigned at the order rank. These reads contributed more than 25% of the total reads in some samples.

**Figure 2.**
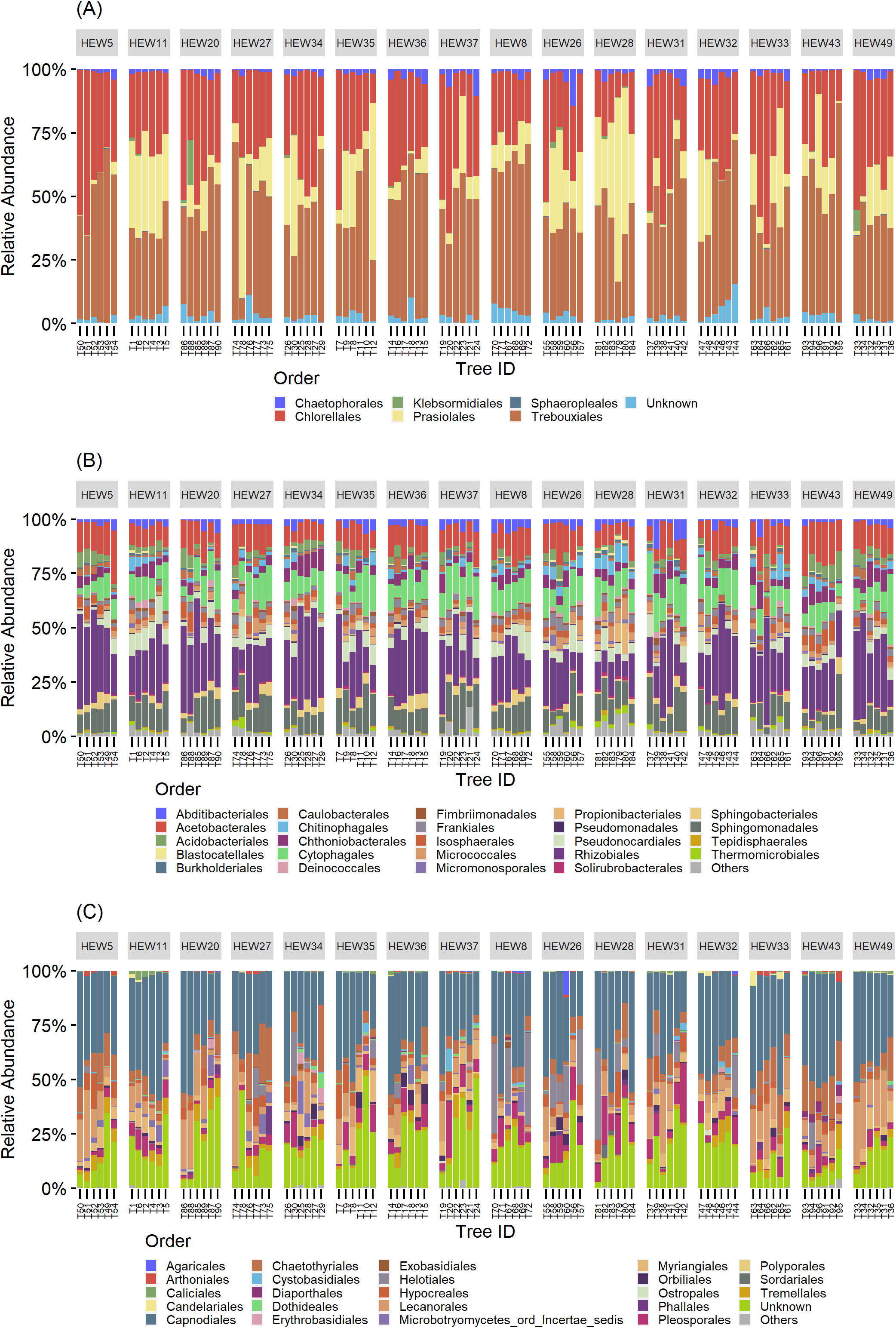
Community bar charts showing the relative abundance of the 25 most abundant orders split by sampling plots. From left to right: the first eight plots are under a low-, and the next eight under a high-management regime. Bars within plots represent individual trees, from large to small tree-size class (2 trees each), from left to right.

### Intra-group interactions

We inferred ASV interaction networks for all three microbial groups (Figure 3). Each ASV entering the networks was present in at least 1% of the samples resulting in 129 ASVs for the algae, 628 for bacteria and 289 for fungi. The algal network (Figure 3 (A)) had a diameter of ten (i.e., the longest shortest path between two nodes was through ten edges), an average path length of four and a modularity score of 0.575. Modularity scores > 0.4 indicate strong modularity ((Newman, 2006).

**Figure 3.**
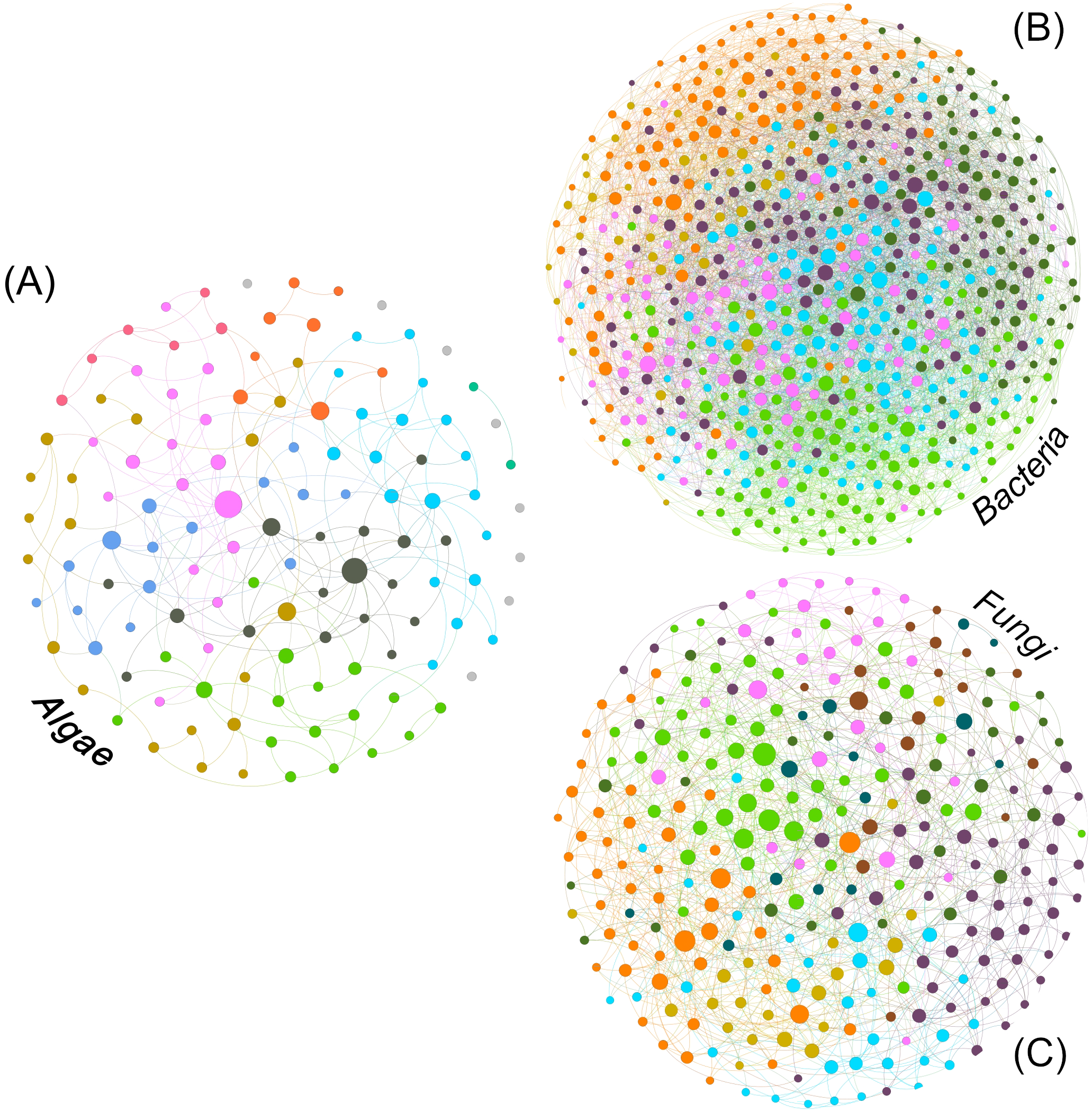
Intra-group ASV interaction network for algal (A), bacterial (B) and fungal (C) fractions of the bark microbiome. The size of the nodes is proportional to the value of betweenness centrality (based on Brandes, 2001) and colours correspond to modules (Blondel et al., 2008).

The diameter for the fungal network (Figure 3 (C)) was 7, with an average path length of ∼3.2 and a modularity slightly lower than the algae at 0.434. The bacterial network (Figure 3 (B)) was denser and more interconnected with a diameter of 5, an average path length of ∼2.7 and a modularity of 0.335.

The algal network could be subdivided into 9 different modules, 5 of which consisting of more than 10 ASVs (see table 3). There were also 8 ASVs that did not interact with any other taxon in the network. The algal module with the highest number of nodes was module 2 (gold colour, Figure 3 (A)) with the most abundant ASV belonging to the genus *Desmococcus* (relative abundance = ∼47% in the module, Table 3).

**Table 3.** Taxonomic description of the most abundant ASVs and their relative abundance for modules with more than 10 ASVs.

| ASV ID | # | Kingdom | Phylum | Class | Order | Family | Genus | Rel. abund. |
| --- | --- | --- | --- | --- | --- | --- | --- | --- |
| **Algae** |  |  |  |  |  |  |  |  |
| ASV_4 | 2 | Eukaryota | Chlorophyta | Trebouxiophyceae | Prasiolales | Prasiolaceae | Desmococcus | 0.47 |
| ASV_1 | 4 | Eukaryota | Chlorophyta | Trebouxiophyceae | Trebouxiales | Trebouxiaceae | Symbiochloris | 0.8 |
| ASV_7 | 5 | Eukaryota | Chlorophyta | Trebouxiophyceae | Trebouxiales | Trebouxiaceae | Trebouxia | 0.4 |
| ASV_2 | 7 | Eukaryota | Chlorophyta | Trebouxiophyceae | Chlorellales | Chlorellaceae | Apatococcus | 0.38 |
| ASV_33 | 9 | Eukaryota | Chlorophyta | Chlorophyceae | Chaetophorales | Chaetophoraceae | Diplosphaera | 0.44 |
| **Bacteria** |  |  |  |  |  |  |  |  |
| ASV_2 | 0 | Bacteria | Proteobacteria | Alphaproteobacteria | Acetobacterales | Acetobacteraceae | Acidiphilium | 0.47 |
| ASV_8 | 1 | Bacteria | Proteobacteria | Alphaproteobacteria | Rhizobiales | Beijerinckiaceae | Methylocella | 0.13 |
| ASV_210 | 2 | Bacteria | Abditibacteriota | Abditibacteria | Abditibacteriales | Abditibacteriaceae | Abditibacterium | 0.08 |
| ASV_54 | 3 | Bacteria | Proteobacteria | Alphaproteobacteria | Acetobacterales | Acetobacteraceae | Acidiphilium | 0.11 |
| ASV_38 | 4 | Bacteria | Proteobacteria | Alphaproteobacteria | Acetobacterales | Acetobacteraceae | Acidiphilium | 0.11 |
| ASV_6 | 5 | Bacteria | Proteobacteria | Alphaproteobacteria | Rhizobiales | Beijerinckiaceae | 1174-901-12 | 0.15 |
| ASV_46 | 6 | Bacteria | Abditibacteriota | Abditibacteria | Abditibacteriales | Abditibacteriaceae | Abditibacterium | 0.13 |
| **Fungi** |  |  |  |  |  |  |  |  |
| ASV_11 | 0 | Fungi | Ascomycota | Lecanoromycetes | Lecanorales | Lecanoraceae | Scoliciosporum | 0.19 |
| ASV_23 | 1 | Fungi | Ascomycota | Eurotiomycetes | Chaetothyriales | Herpotrichiellaceae | Capronia | 0.58 |
| ASV_51 | 2 | Fungi | Ascomycota | Dothideomycetes | Capnodiales |  |  | 0.14 |
| ASV_32 | 3 | Fungi | Ascomycota | Dothideomycetes | Capnodiales | Mycosphaerellaceae |  | 0.25 |
| ASV_43 | 4 | Fungi | Ascomycota | Dothideomycetes | Capnodiales |  |  | 0.39 |
| ASV_4 | 5 | Fungi | Ascomycota |  |  |  |  | 0.36 |
| ASV_6 | 6 | Fungi | Ascomycota | Leotiomycetes | Helotiales |  |  | 0.3 |
| ASV_1 | 7 | Fungi | Ascomycota | Dothideomycetes | Capnodiales |  |  | 0.88 |
| ASV_61 | 8 | Fungi | Ascomycota |  |  |  |  | 0.18 |

The bacterial network consisted of 7 modules, all of which with more than 10 ASVs and no taxa that have no connections. In this case, module 6 (purple colour in Figure 3 (B)) was the module with the highest count of taxa, with an ASV assigned to the genus *Abditibacterium* being the predominant strain (12% of reads in the module, Table 3).

For fungi, the network clustered into 9 different modules, all containing more than ten ASVs and no unconnected taxa. Also in this case, module two (purple colour, Figure 3 (C)) was the module with the highest number of taxa for the fungal network, with an ASV belonging to the order Capnodiales – not assignable to a higher taxonomic rank (Table 3) - with the highest relative abundance in the module (14%).

Network structure was examined by identifying nodes with the highest number of shortest path going through them (betweenness centrality), indicating taxa that are important for the connectivity of the network. We defined so called hub taxa as the five taxa with the highest betweenness centrality (Table 4). In the algal network these hub taxa belonged to two orders, Chlorellales and Trebouxiales, and three different genera. Three ASVs were assigned to the genus *Apatoccocus*, and one to *Trebouxia* and *Symbiochloris*, respectively. Bacterial hub taxa showed a higher diversity than the algae with hub taxa belonging to 5 different orders. The genera include *Tundrisphaera, Hymenobacter, Actinomycetospora* and *Oligoflexus*. One of the ASVs was assigned to the group 1174-901-12, a group of uncultured bacterial strains within the order Rhizobiales. Many of the fungal hub taxa were not assigned at the genus rank, with the exception of two ASVs belonging to the genera *Tremella* and *Aureobasidium*. Two more ASVs were assigned to the order Capnodiales while one was only assigned at the phylum rank.

**Table 4.** Taxonomic information for the hub taxa of the respective network, identified based on their betweenness centrality.

| ASV ID | Kingdom | Phylum | Class | Order | Family | Genus |
| --- | --- | --- | --- | --- | --- | --- |
| **Algae** |  |  |  |  |  |  |
| ASV 10 | Eukaryota | Chlorophyta | Trebouxiophyceae | Chlorellales | Chlorellaceae | Apatococcus |
| ASV 11 | Eukaryota | Chlorophyta | Trebouxiophyceae | Chlorellales | Chlorellaceae | Apatococcus |
| ASV 13 | Eukaryota | Chlorophyta | Trebouxiophyceae | Trebouxiales | Trebouxiaceae | Symbiochloris |
| ASV 22 | Eukaryota | Chlorophyta | Trebouxiophyceae | Trebouxiales | Trebouxiaceae | Trebouxia |
| ASV 8 | Eukaryota | Chlorophyta | Trebouxiophyceae | Chlorellales | Chlorellaceae | Apatococcus |
| **Bacteria** |  |  |  |  |  |  |
| ASV 162 | Bacteria | Planctomycetota | Planctomycetes | Isosphaerales | Isosphaeraceae | Tundrisphaera |
| ASV 19 | Bacteria | Proteobacteria | Alphaproteobacteria | Rhizobiales | Beijerinckiaceae | 1174-901-12 |
| ASV 559 | Bacteria | Bacteroidota | Bacteroidia | Cytophagales | Hymenobacteraceae | Hymenobacter |
| ASV 79 | Bacteria | Actinobacteriota | Actinobacteria | Pseudonocardiales | Pseudonocardiaceae | Actinomycetospora |
| ASV 855 | Bacteria | Bdellovibrionota | Oligoflexia | Oligoflexales | Oligoflexaceae | Oligoflexus |
| **Fungi** |  |  |  |  |  |  |
| ASV 16 | Fungi | Ascomycota | Dothideomycetes | Capnodiales |  |  |
| ASV 18 | Fungi | Basidiomycota | Tremellomycetes | Tremellales | Tremellaceae | Tremella |
| ASV 4 | Fungi | Ascomycota |  |  |  |  |
| ASV 56 | Fungi | Ascomycota | Dothideomycetes | Dothideales | Aureobasidiaceae | Aureobasidium |
| ASV 65 | Fungi | Ascomycota | Dothideomycetes | Capnodiales |  |  |
| **Combined** |  |  |  |  |  |  |
| ASV 116 | Fungi | Ascomycota | Dothideomycetes | Pleosporales | Massarinaceae | Massarina |
| ASV 65 | Fungi | Ascomycota | Dothideomycetes | Capnodiales |  |  |
| ASV 116 | Bacteria | Verrucomicrobiota | Verrucomicrobiae | Chthoniobacterales | Chthoniobacteraceae | LD29 |
| ASV 713 | Bacteria | Bacteroidota | Bacteroidia | Chitinophagales | Chitinophagaceae | Edaphobaculum |
| ASV 105 | Eukaryota | Chlorophyta | Trebouxiophyceae | Trebouxiales | Trebouxiaceae | Trebouxia |

### Inter-group interactions

The combined network of algae, bacteria and fungi (Figure 4) displayed a diameter of 4, an average path length of ∼2.6 and a modularity of 0.259, making this network more densely connected than the bacterial network. Out of the eight modules, modules two (29%) and three (26%) accounted for more than half of the total reads.

**Figure 4:**
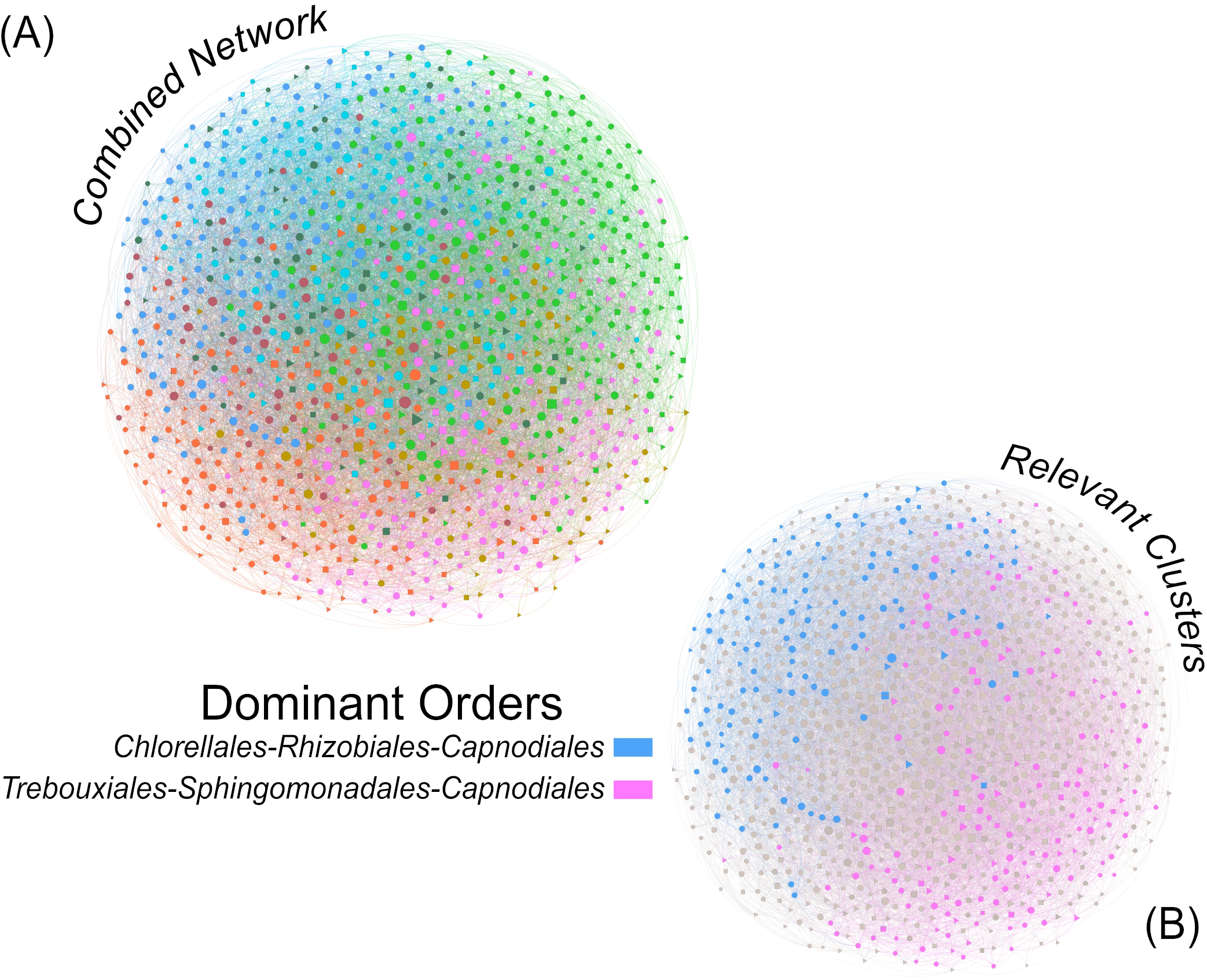
Inter-group ASV interaction network integrating algal, bacterial and fungal taxa for a description of the interactions within the complete bark microbiome (A). The two coloured modules (B) are the two most abundant modules of the whole dataset, accounting for 29% (pink) and 26% (blue) of the total reads. The size of the nodes is proportional to the value of betweenness centrality (based on Brandes, 2001) and colours correspond to modules (Blondel et al., 2008).

The top orders for algae, bacteria and fungi in the relatively most abundant module (module 2, pink colour in Figure 4 (B)) were Trebouxiales, Sphingomonadales and Capnodiales, respectively. Algae and fungi accounted for up to 36 and 26% of the module reads, respectively, while Sphingomonadales only contributed 5% of the reads. Trebouxiales contributed 87% of the algal reads, Sphingomonadales 30% of bacterial reads, and Capnodiales 77% of the fungal reads. In total, algae contributed 42%, bacteria 19%, and fungi 34% of the reads assigned to module 2.

In the second most abundant module (module 3, blue colour in Figure 4 (B)) Chlorellales were the most important algal order, Rhizobiales the most important bacterial order, while the most important fungal order was again Capnodiales. Chlorellales contributed 18%, Rhizobiales 16% and Capnodiales only 5% of the module reads. Chlorellales accounted for 43% of the algal reads, Rhizobiales 48% of the bacterial reads, and Capnodiales 34% of the fungal reads. Overall, module three consisted of 42% algae, 34% bacteria and 10% fungi.

The hub taxa of the combined network (Table 4) contained members of all t three microbial groups, specifically one alga, two bacteria and two fungi. Four ASVs could be assigned down to genus rank, while one fungal ASV – a hub taxon present also in the fungal network – could only be assigned to order rank (Capnodiales). The other fungus belonged to the genus *Massarina*, while the two bacteria ASVs belonged to the genus *Edaphobaculum* and the uncultured group LD29 within the order Chthoniobacterales. The only algal taxon represented the common lichen-forming genus *Trebouxia*.

### Drivers of changes in community composition

To investigate differences in community composition we plotted the results of the PCA (Figure 5). The differences between the high and low management intensity are not readily visible by looking at the clusters of the two groups. Yet, there are significant differences between the groups as revealed by the PERMANOVA results (algae: p < 0.05, bacteria: p < 0.01, fungi: p < 0.001). The dispersion permutation test was significant (algae: p < 0.05, bacteria: p < 0.05, fungi: p < 0.05) indicating a heterogeneous dispersion within groups. If on one hand this might suggest that the results are not reliable, Anderson and Walsh (2013) showed that PERMANOVA is not sensitive against heterogeneity, compared to other analyses. Management intensity explained only ∼2% of the variance in the dataset, suggesting a subtle yet significant effect.

**Figure 5.**
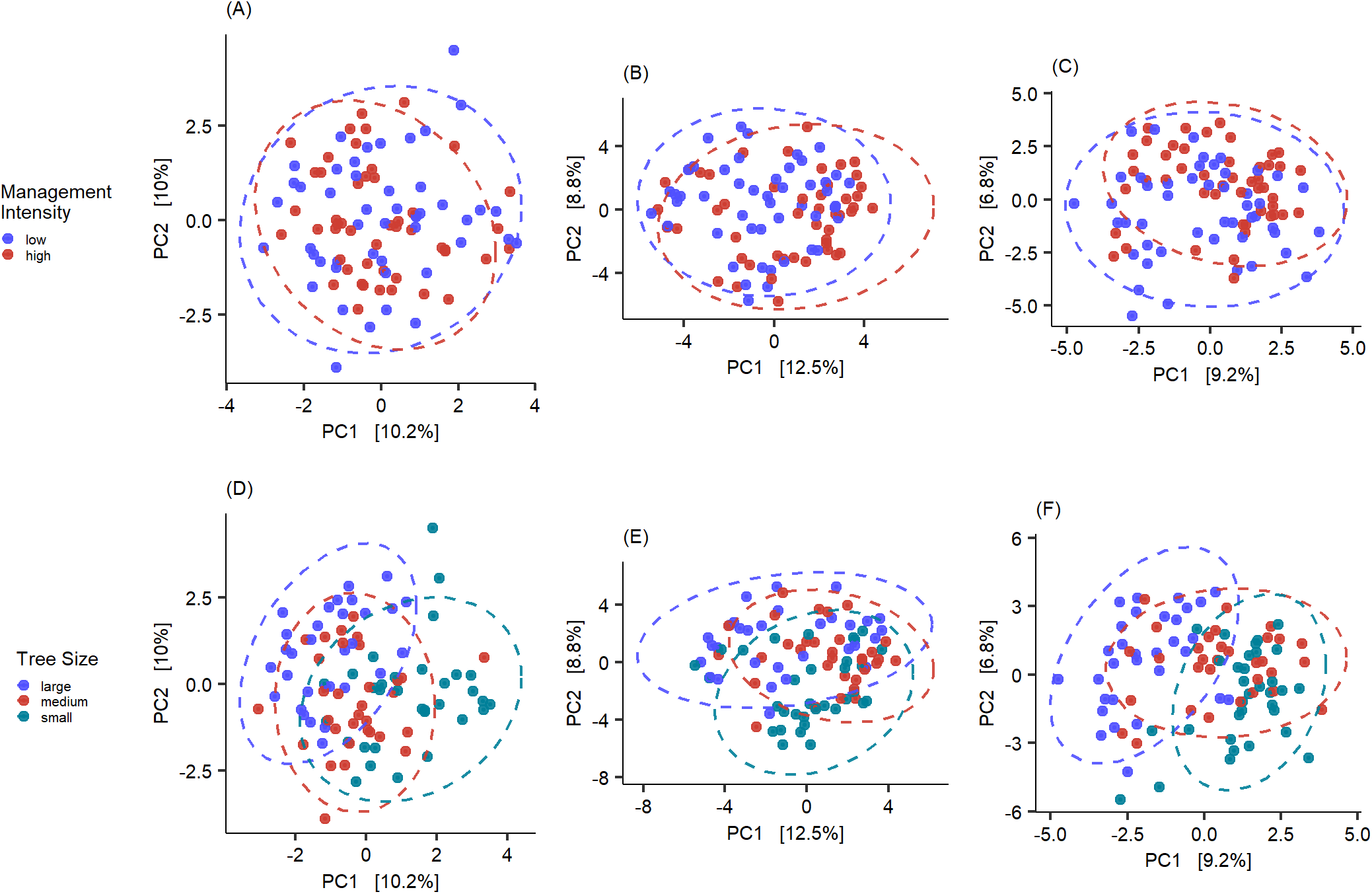
Overview of the Principle Component Analysis for algae (A and D), bacteria (B and E) and fungi (C and F). Colours indicate the tested groups and axes are scaled to the proportion of variance explained by the displayed Principal Component (Nguyen and Holmes, 2019).

The tree sizes clustered in a much clearer structure, with less overlap of the 95% confidence interval ellipses. The PERMANOVA analysis confirmed this observation, with highly significant results for all microbial groups (p < 0.001 for all), while the permutation test indicated homogenous dispersion within the groups (p > 0.05).

Further examination of the community structure showed that the differences in community composition were also visible through significantly differentially abundant taxa (Supplementary Table 2). The difference between management intensities was little, with only four differentially abundant ASVs. The tree-size classes, however, showed a different signal and confirmed the stronger differences indicated by the PERMANOVA results. Between large and medium trees we found nine algae, four bacteria and ten fungal taxa that showed significant abundance differences. A larger difference in abundances could be seen between large and small trees, with 19 algae, 45 bacteria and 33 fungi that were differentially abundant. Almost no difference could be observed between communities of medium and small trees with only two algal and one bacterial ASVs with significant changes in abundance.

## Discussion

In this study we used a multi-kingdom metabarcoding approach to investigate the microbiome (algae, bacteria, fungi) on the bark of beech from sites with different forest management intensity in the Hainich-Dün region, Thuringia, Germany. We provide a first characterization of aboveground bark-associated microbial communities in beech forests, as well as an account of microbial inter-kingdom interactions. Additionally, by testing how community diversity and structure of the different organismal groups change in relation to land-use intensity and tree size, we identify potential drivers of community assembly and provide sampling recommendations for studying the bark microbiome at broader spatial scales.

### Diversity of the bark-associated beech microbiome

Our results show that the bark of beech harbours highly diverse algal, bacterial and fungal communities.

Algal diversity is mainly represented by species of the families Trebouxiophyceae and Chlorophyceae, in particular by genera commonly found in subaerial environments and already detected in forest settings (Štifterová and Neustupa (2015)). Algae of the genus *Apatococcus*, a “flagship” taxon of above-ground ecosystems (Rindi 2007), are among the most abundant and interconnected members of the algal microbiome. Another abundant alga is *Desmococcus* sp., which typically forms visible powdery, greenish layers on the bark of trees in association with *Apatococcus* and other subaerial green algae (Rindi, 2007). Both are part of what is possibly the most tolerant subaerial algal community, being able to thrive even in urban areas (Barkman, 1958). Furthermore, our results confirm *Symbiochloris* as a common player in the dermo- and phyllosphere (Škaloud et al., 2016; Zhu et al., 2018) as well as other Trebouxiales, an order consisting of free-living as well as lichen-forming green algae (Sanders and Masumoto, 2021).

Compared to the algae and fungi, the bacterial community is far more diverse. Similar as in Aschenbrenner et al., (2017), the bacterial community is dominated by the class Alphaproteobacteria, in particular Rhizobiales and Acetobacteriales. Among the Rhizobiales we detected *Methylocella* sp., a facultative methanotroph adapted to various nutrients (acetate, pyruvate, succinate, malate, and ethanol) (Dedysh and Dunfield, 2011), and 1174-901-12, previously described as an early colonizer of aerial environments (Romani et al., 2019) and a known member of the phyllosphere (Ares et al., 2021). In the case of Acetobacterales, *Acidiphilium* is among the most abundant taxa. The genus consists of aerobic bacteria with photosynthetic pigments, with a pH range that matches well to the pH of beech bark (∼4.4 in Asplund et al., 2015) and that do not overlap in metabolic demands with *Methylocella* (Hiraishi and Imhoff, 2015).

Another abundant class are the *Abditibacteria*, especially the genus *Abditibacterium* whose representatives are well adapted to low-nutrient conditions and have already been reported on tree bark (Tahon et al., 2018; Kobayashi and Aoyagi, 2019). Among the bacteria hub taxa we find genera from classes already reported for tree-associated microbiomes, such as *Oligoflexus* in bryophytes (Ma et al., 2017), *Actinomycetospora* and 1174-901-12 on tree bark and lichens ((Yamamura et al., 2011), (Ares et al., 2021)), *Hymenobacter*, a desiccation- and radiation-tolerant bacterium often found in soil (Buczolits and Busse, 2015), and *Tundrisphaera*, previously only isolated from lichen-dominated tundra soils (Kulichevskaya et al., 2017). Contrarily to Aschenbrenner et al., (2017), we did not find major contributions from the genera *Burkholderia* and *Pseudomonas*, possibly indicating that these genera are specific to the sycamore maple (*Acer pseudoplatanus*) bark.

The investigation of fungal diversity was somewhat hindered by low taxonomic resolution with many ASVs that could not be assigned past phylum rank. This may result from a lack of resolution in public fungal databases combined with the presence of several unknown taxa in our dataset. Abundant members of the beech bark fungal community are the so-called black yeasts, e.g., *Capronia* and *Aureobasidium*, which are known to occur on tree bark and leafs (Untereiner and Malloch, 1999; Andrews et al., 2002), but also decaying wood (Cooke, 1959) and on other fungi or lichens as secondary saprobionts (Untereiner and Malloch, 1999). Other common fungi belong to the genus *Tremella*, known mycoparasites (Zugmaier et al., 1994). Among the lichen-forming fungi, the most common fungus in our dataset belongs to the genus *Scoliciosporum*, a genus of crustose lichens that was already reported on beech bark (e.g., Dymytrova, 2011). The biggest contributors at the order rank are members of the Capnodiales (Dothideomycetes) and Helotiales (Leotiomycetes), whose species have been shown to associate with the lichen microbiome (Suija et al., 2014; Smith et al., 2020). Yet, more research into these orders is needed as they are taxonomically and ecologically highly diverse and include a large diversity of life forms, from lichenized, to mycoparasytic, epi-, ecto-, endophytic, as well as saprobiontic species (Tedersoo et al., 2009; Abdollahzadeh et al., 2020). In conclusion, more research is needed in order to confirm the role of the bark habitat as a reservoir of novel fungal diversity. This could possibly be done by combining genetics with culture-based approaches.

### Biotic interactions and inter-kingdom synergies in the bark microbiome

The higher modularity scores of the fungal and especially algal networks may indicate higher specialization or niche differentiation in these groups (Augustyn et al., 2016). In contrast, bacteria are less clearly divided into ecological modules, which potentially indicates closer interactions between all taxa as there seems to be no split into specialized groups. Further analyses based on a broader dataset are needed to exclude that the observed patterns are an artefact of the overall higher diversity found in bacteria.

The results from the combined, inter-kingdom co-occurrence analysis indicate that algal and fungal specialists might be connected through a common set of bacteria. It is tempting to speculate that the interactions between Rhizobiales and Chlorellales (mostly represented by members of the genus *Apatococcus*) observed in the main ecological cluster in our dataset are of symbiotic nature, as Rhizobiales are well-known beneficial partners in plant-microbe interactions and common associates of lichens (Erlacher et al., 2015; Grube et al., 2015). Positive interactions among Sphingomonadales, Trebouxiales, and Capnodiales – all known occupants of bark substrates – characterize the secondly most important cluster. The bacterial genus *Sphingomonas* is very common in above-ground forests habitats (Vorholt, 2012), exhibiting facultative photosynthesis.

Finally, we identified the most highly connected taxa (hubs), i.e. taxa that are crucial for the stability of the ecological network (Banerjee et al., 2018). For bacteria, the hub taxa belong to the genera LD29 (Verrucomicrobiota) and *Edaphobaculum* (Bacteroidetes). Little is known about their ecology, with LD29 particularly abundant in lichen thalli (Aschenbrenner et al., 2017), and *Edaphobaculum* previously found in soils where it contributes to the creation of biofilms (Keuschnig et al., 2021). As for the fungi, one of the hub taxa belongs to the genus *Massarina* (Pleosporales, Dothideomycetes), a genus which is seemingly common on *Fagus sylvatica* (Zhang et al., 2009). Interestingly, for the algae, the hub taxon is a member of the genus *Trebouxia*, the most common lichen-forming alga.

### Bark microbiome responds to tree size, but not to intensity of forest management

The intensity of the forest management regime has virtually no effect on microbial community diversity and structure in our study area. This might be a result of a forest management plan that avoids clear cuts and carefully selects trees to harvest, which leads to a uniform forest structure in the study area (Schall et al., 2020). Based on a broader sampling including this and other two large forest areas in Germany, Boch et al., (2021) showed that an increase in forest management intensity is linked to reduced lichen species richness. A larger sampling effort covering a broader gradient of land-use intensity is therefore required to test whether the bark-associated microbiome differs in response compared to the macroepiphytes.

We found significant differences in diversity and composition of the bark microbiome according to different tree-size classes. The lower microbial diversity found on larger (older) trees for algae and fungi is probably the result of environmental filtering on highly heterogeneous pioneer communities over time. This is particularly evident when comparing large and small trees, thus suggesting slow succession of these microbiomes toward final community composition. Lastly, results from the spatial autocorrelation analysis underpin random assembly of the microbial bark community at the local (plot) level, with a high heterogeneity between trees.

### Conclusions and sampling recommendations

In this pioneering study we provide novel insights into the diversity, spatial context, and biotic interactions that characterize the beech bark microbiome in Central European forests. We showed that there are predictable community shifts depending on tree age. These represent the first steps towards proposing a framework of community assembly on forest tree bark, a ubiquitous, ecologically relevant, yet overlooked component of terrestrial habitats.

Taken together, our results show that a single tree does not adequately characterize the bark-associated microbial community at plot level. To capture most of the microbial diversity, we recommend sampling using a spatially random approach with a balanced representation of the main tree-size classes present in the plot. Samples taken from multiple trees can then be combined into a composite sample. The use of composite samples ensures relatively low costs for obtaining adequate sequencing depths while maximizing the spatial range of the study and the number of plots, allowing for easy upscaling to large areas and/or environmental gradients.

## Acknowledgements

We thank the managers of the Hainich-Dün Exploratory, Anna K. Franke, and all former managers for their work in maintaining the plot and project infrastructure; Victoria Grießmeier for giving support through the central office, Andreas Ostrowski for managing the central data base, and Markus Fischer, Eduard Linsenmair, Dominik Hessenmöller, Daniel Prati, Ingo Schöning, François Buscot, Ernst-Detlef Schulze, Wolfgang W. Weisser and the late Elisabeth Kalko for their role in setting up the Biodiversity Exploratories project. We thank the administration of the Hainich national park, as well as all landowners for the excellent collaboration. The work has been funded by the DFG Priority Program 1374 “Biodiversity-Exploratories” (SCHM 1711/8-1 and GR 5437/4-1). Field work permits were issued by the responsible state environmental offices of Thüringen. Additionally, we thank Ulrich Pruschitzki for his valuable help in the forest and Dr. Jürgen Otte for helpful comments on laboratory procedures and technical assistance. This work has been deposited on bioRxiv as a preprint (DOI: ….).

## Data availability statement

All data necessary to replicate the analyses is publicly available in the repository of the Biodiversity Exploratories BExIS (https://www.bexis.uni-jena.de/). Accession numbers: 31183, 31185, 31186 (taxonomy tables); 31160, 31161, 31162 (ASV tables); 31157, 31158, 31159 (DNA concentrations for use with *decontam*), 31163, 31164, 31165, 31166 (modules obtained from *Gephi*) and 31167 (metadata table). Raw reads of the samples are available at Genbank: Accession number XXXX. Sequences attributed to the ASVs are available at NCBI SRA under accession number XXXX. The code for the processing of the raw reads, sample inference and all analyses is available from Github at https://github.com/LukDrey/beech_micro_communities.

[All this information will be made public after acceptance of the manuscript and is made available to the reviewers as supplementary data.]

